# Cryo-EM structure of influenza polymerase bound to the cRNA promoter provides insights into the mechanism of virus replication

**DOI:** 10.1101/2025.09.23.677976

**Authors:** Yixi Wu, Minke Li, Huanhuan Li, Tianli sun, Yifan Bai, Shaohui Huang, Yingfang Liu, Huanhuan Liang

## Abstract

Influenza virus polymerase (FluPol) synthesizes the complementary RNA (cRNA) and the viral RNA (vRNA) using distinct *de novo* initiation strategies during genome replication, known as internal and terminal initiation, respectively. The *de novo* initiation mechanisms, especially the internal initiation process, which includes a template realignment process, is still not well understood. Here, we present a cryo-electron microscopy structure of H5N1 FluPol bound to the cRNA promoter. In combination with structural analyses and structure-guided mutagenesis studies, we identified several previously unreported structural features of FluPol essential for internal initiation. The B loop adopts an “open” conformation, allowing translocation of the 3′ terminus of cRNA (3′-cRNA) template into the catalytic cavity. The dynamic of incoming 3′-cRNA template is limited by the priming and realignment loop (PR loop), which facilitates the cRNA template realignment process. An asparagine cluster above the catalytic cavity is required for polymerase activity. Our findings provide structural insights into the mechanism of replication internal initiation of FluPol.

## Introduction

Influenza viruses pose a significant threat to human life due to their ability to rapidly evolve and cause seasonal outbreaks as well as occasional pandemics. Influenza viruses are negative-sense, single-stranded RNA viruses with segmented genomes. Their genomes are characterized by conserved 3′ and 5′ termini (known as promoter regions), which tightly bind to the RNA-dependent RNA polymerase (RdRp, also called FluPol). FluPol is responsible for both transcription and replication of the viral genomes. It utilizes the viral RNA (vRNA) as a template to generate capped and polyadenylated transcription products in a primer-dependent manner, while also generating replication products through *de novo* initiation mechanisms^1–4^.

In recent years, numerous high-resolution structures of FluPol have been revealed^4–14^. These structures have elucidated that FluPol is composed of a stable core region, which includes the C-terminal domain of the PA subunit (PA-C), the PB1 subunit, and the N-terminal domain of the PB2 subunit (PB2-N), as well as the highly flexible domains that include the N-terminal domain of the PA subunit (PA-N) and the C-terminal domain of the PB2 subunit (PB2-C). Notably, most resolved structures of FluPol are in transcription states, providing insights into how the FluPol undergoes conformational shifts to facilitate the transcription initiation, elongation and termination processes^4–7^. Previous functional studies have demonstrated that replication of influenza viral genome includes cRNA synthesis and vRNA synthesis processes^15^. FluPol initiates the cRNA intermediate synthesis terminally using the vRNA template, which could be subdivided into three steps. First, 3′ terminus of vRNA (3′-vRNA) template of FluPol translocates and positions the first and second nucleotides (+1U and +2C) at the catalytic sites (referred to as “priming”). Second, the catalytic sites recruit and polymerize the ATP and GTP to generate the dinucleotide pppApG. Third, the catalytic sites continue cRNA synthesis under NTPs incorporation. Subsequently, the cRNA intermediate serves as the template for initiating vRNA synthesis internally employing a unique priming and realignment mechanism^15^, which could be subdivided into five steps. First, 3′-cRNA template of FluPol translocates and positions the fourth and fifth nucleotides (+4U and +5C) at the catalytic site (referred to as “priming”). Second, the catalytic sites recruit and polymerize the ATP and GTP to generate the dinucleotide pppApG. Third, the first and second nucleotides (+1U and +2C) of cRNA template realigns to the active sites (referred to as “realignment”). Fourth, the first and second nucleotides (+1U and +2C) form base pairing with dinucleotide pppApG at the catalytic sites. Fifth, the catalytic sites continue vRNA synthesis under NTPs incorporation^16^.

Despite its functional importance, the structural basis and precise molecular dynamics governing FluPol’s replication process remain poorly understood. Only a limited number of structures in replication states have been reported. For instance, the structure of influenza B virus polymerase (FluPol_B_) bound to the 5′ terminus of the cRNA promoter revealed remarkable rearrangements within the PB2 subunit compared to those in transcription states^14^. Additionally, the structure of influenza D virus polymerase (FluPol_D_) in complex with the cRNA promoter revealed that 3′-cRNA preferentially binds to the secondary binding site on the surface of FluPol in the replication pre-initiation state^11^. These structures demonstrate substantial differences between FluPol’s replication and transcription states. Some structures further suggest host factor involvement in the viral polymerase replication process. For example, the structure of the replication platform, assembled through bridging of two polymerase molecules by the host factor ANP32A, illustrates how the nascent vRNA products are encapsidated^12^. Despite the aforementioned studies, many questions regarding influenza virus polymerase replication remain unresolved. For instance, how FluPol facilitates this translocation and stabilizes the 3′-RNA within the cavity remains elusive. Additionally, even in cell-free biochemical assays lacking host factors, the FluPol can still efficiently execute both terminal and internal replication mechanisms. In other words, how does the FluPol autonomously regulate the terminal and internal initiation processes? Compared to the terminal initiation mechanism, the internal mechanism remains less well understood due to its inherent complexity, particularly how FluPol achieves realignment during internal initiation. The key structural elements or amino acids involved in these processes are poorly characterized.

Previous mutagenesis screening and *in vitro* replication assays have identified several structural elements and amino acids that affect replication initiation of FluPol. For instance, the priming loop, which extends from the thumb domain, serves as a stacking platform for NTP incorporation during *de novo* initiation^17^. Additionally, we have previously reported that the motif I on the PB1 subunit plays an important role in stabilizing the priming loop, thereby facilitating the replication initiation process^18^. Furthermore, it was reported that the V273A mutation in the helix-loop-helix motif (residues 269-282) led to an increase in production of failed realignment products^19^. The mechanism by which Val273 regulates the cRNA template realignment process remains unclear.

Notably, this template realignment mechanism is also employed in other viral polymerases, such as those of the La Crosse virus (LACV), Hantaan virus, arenaviruses, and rotaviruses^20–23^, in which several structural elements involved in replication initiation have been identified. Previous reports have shown that the motif B (B loop), a typical motif in the right-hand RdRp structure, corrects the position of the RNA template by allosteric regulation^24^. The B loop can adopt either an “open” or “closed” conformation. It allows the template RNA to enter the active site in the “open” conformation, while it obstructs the pathway of the RNA template to the active site of FluPol in the “closed” conformation^24^. Additionally, the PR loop, located above the catalytic site in the polymerase of LACV, is involved in stabilizing the 3′ terminus of the template^23,25–27^. In the structure of LACV polymerase, the residues Met989, Ile990 and Ser991 in the PR loop interact with the 3′ terminus of the RNA template, facilitating the replication initiation^23^. We speculate that these structural elements play similar roles in FluPols, but this hypothesis lacks support from previous structural and functional studies.

In this study, we present a structure of avian influenza A/Goose/Guangdong/3/1997 (H5N1) virus polymerase bound to the cRNA promoter (FluPol_H5N1_-cRNA). This structure revealed several structural elements that regulate the replication initiation of FluPol. This study advances our understanding of the replication internal initiation mechanism of FluPol.

## Results

### The FluPol_H5N1_-cRNA structure represents a pre-catalytic state

To better understand the replication internal initiation mechanism of FluPol, we determined a structure of influenza H5N1 polymerase bound to the cRNA promoter at an average resolution of 2.98 Å (Fourier shell correlation at 0.143 criterion) using cryo-electron microscopy (cryo-EM) three-dimensional reconstruction methods (Table 1 and Fig. S1). This structure contains a stable core region consisting of PA-C (residues 201- 716), PB1 (residues 1-631, 661-668) and PB2-N (residues 61-82, 90-103) but lacks the flexible PA-N and PB2-C domains (Fig. 1a-b). The PA-C is located at the bottom of the FluPol, and the PB1 subunit folds into a typical right-handed RNA-dependent RNA polymerase structure. The 5′ terminus of cRNA (5′-cRNA, 5′- AGCAAAAGCAGGGUGUUU-3′) is fully observed. Distinct from the reported replication pre-initiation state, which is characterized by 3′-cRNA binding to a secondary site on the FluPol surface^11^, the 3′-cRNA in our structure (5′- AACACCCUU-3′) is located at the template entrance channel (Fig. 1c). This observation indicates that the FluPol_H5N1_-cRNA structure probably represents a state ready for replication initiation. In comparison to FluPols in transcription and intermediate states^4,5,14^, we identified several structural features in the FluPol_H5N1_- cRNA structure that are crucial for replication initiation.

**Fig. 1.**
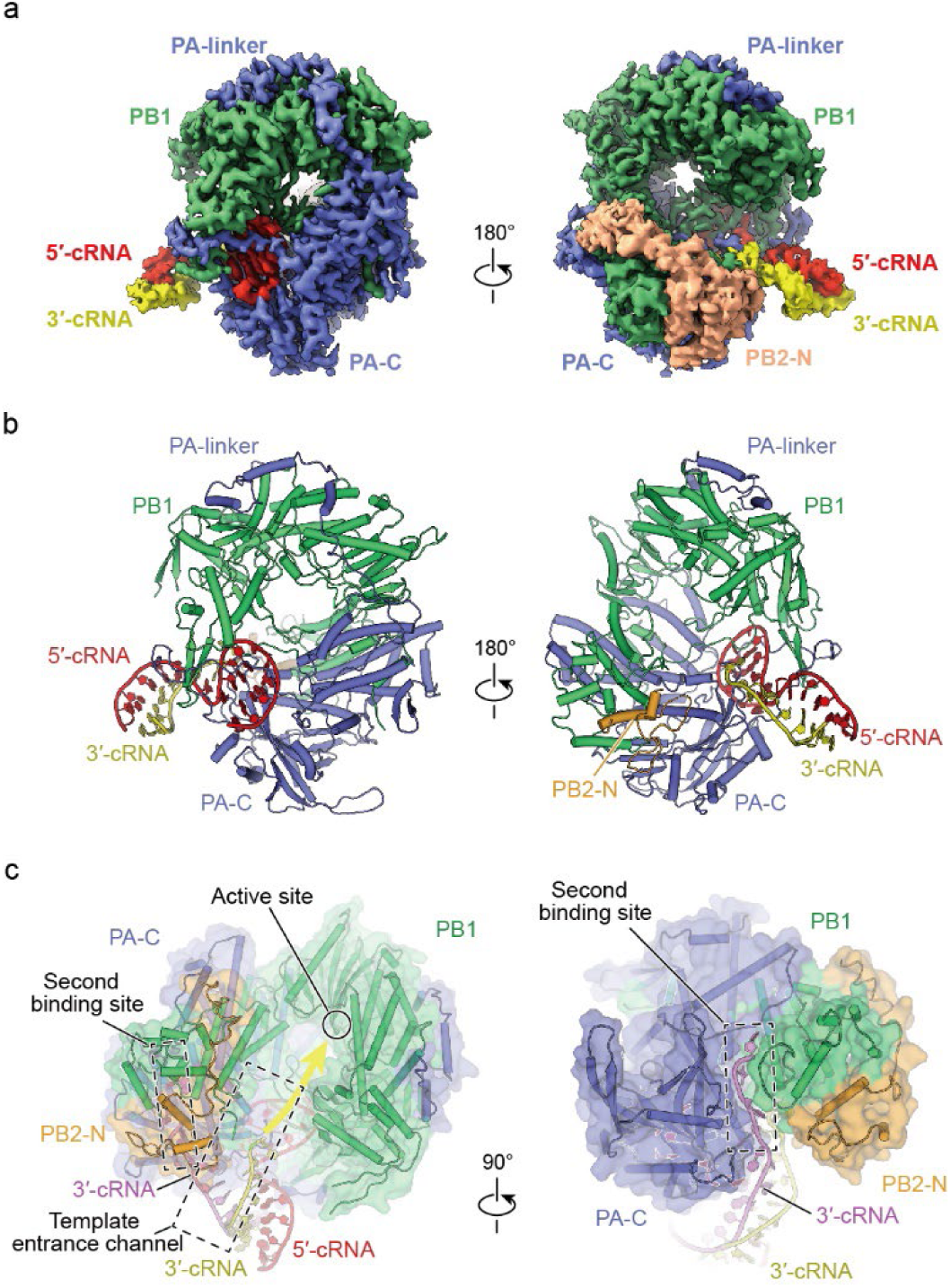
Overall structure of FluPol_H5N1_ bound to the cRNA promoter. **a, b.** Cryo-EM map (**a**) and cartoon representation (**b**) of the FluPol_H5N1_-cRNA structure. The PA, PB1, PB2 subunits, 3′-cRNA and 5′-cRNA promoters are shown in blue, green, orange, yellow and red, respectively. **c.** The partially resolved 3′-cRNA is located at the template entrance channel. The yellow arrow indicates that the 3′-cRNA directs towards the catalytic cavity in the FluPol_H5N1_-cRNA structure (**left)**. This arrangement is notably distinct from the 3′-cRNA (violet) superposed from the FluPol_D_-cRNA structure (PDB 6ABE), which binds to the secondary binding site (**right**).

**Table 1.**
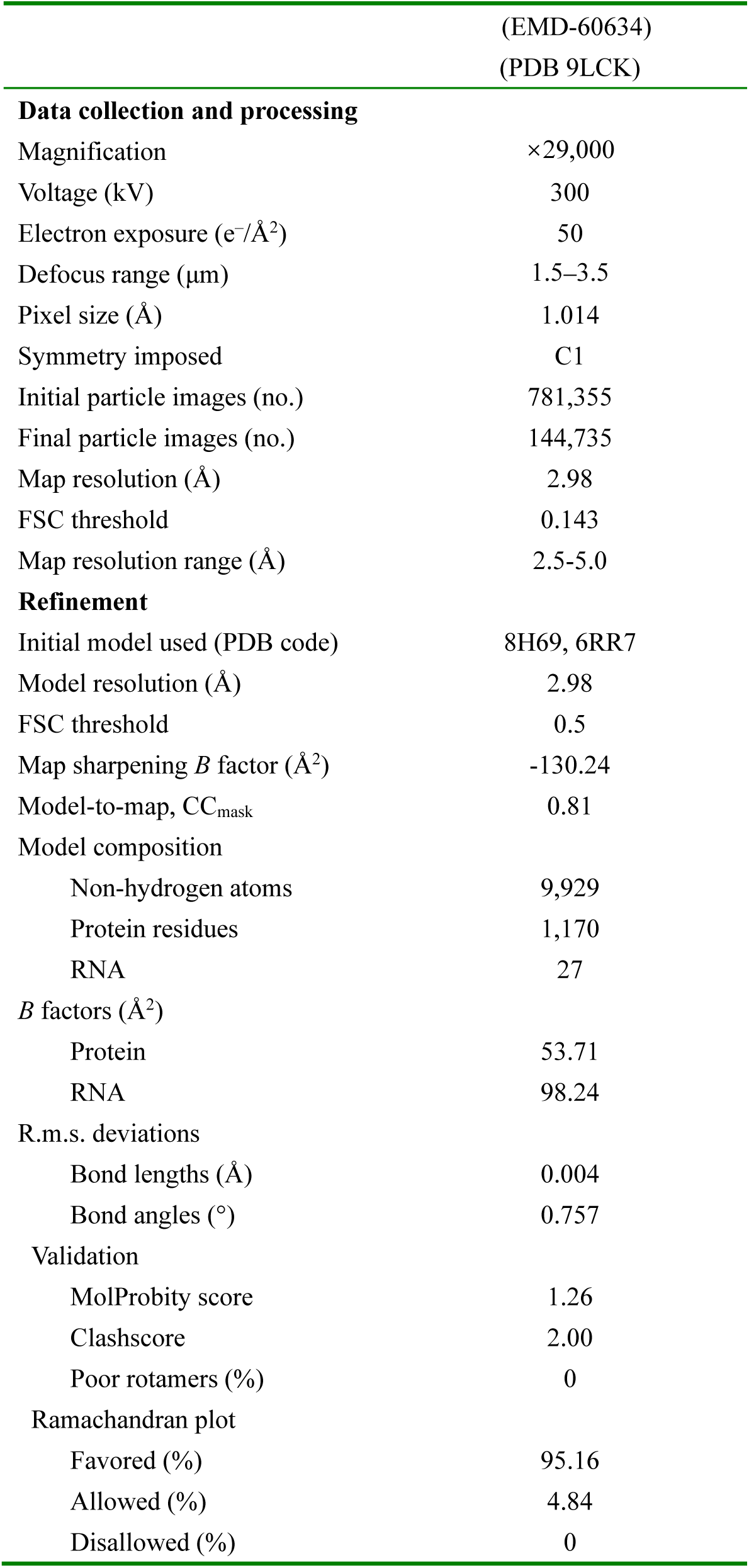
Cryo-EM data collection, refinement and validation statistics.

|  | (EMD-60634) |
| --- | --- |
|  | (PDB 9LCK) |
| **Data collection and processing** |  |
| Magnification | ×29,000 |
| Voltage (kV) | 300 |
| Electron exposure (e<sup>-</sup>/Å<sup>2</sup>) | 50 |
| Defocus range (μm) | 1.5–3.5 |
| Pixel size (Å) | 1.014 |
| Symmetry imposed | C1 |
| Initial particle images (no.) | 781,355 |
| Final particle images (no.) | 144,735 |
| Map resolution (Å) | 2.98 |
| FSC threshold | 0.143 |
| Map resolution range (Å) | 2.5–5.0 |
| **Refinement** |  |
| Initial model used (PDB code) | 8H69, 6RR7 |
| Model resolution (Å) | 2.98 |
| FSC threshold | 0.5 |
| Map sharpening *B* factor (Å<sup>2</sup>) | -130.24 |
| Model-to-map, CC<sub>mask</sub> | 0.81 |
| Model composition |  |
| Non-hydrogen atoms | 9,929 |
| Protein residues | 1,170 |
| RNA | 27 |
| *B* factors (Å<sup>2</sup>) |  |
| Protein | 53.71 |
| RNA | 98.24 |
| R.m.s. deviations |  |
| Bond lengths (Å) | 0.004 |
| Bond angles (°) | 0.757 |
| Validation |  |
| MolProbity score | 1.26 |
| Clashscore | 2.00 |
| Poor rotamers (%) | 0 |
| Ramachandran plot |  |
| Favored (%) | 95.16 |
| Allowed (%) | 4.84 |
| Disallowed (%) | 0 |

### The B loop adopts an “open” conformation required for incoming 3′-RNA template

In the FluPol_H5N1_-cRNA structure, we observed that the B loop on the finger domain retracts from the active site, adopting an “open” conformation—a structural feature also presented in transcription initiation state (Fig. 2a-b). In contrast, the B loop protrudes toward the catalytic cavity in transcription pre-initiation and intermediate state (FluPol_H5N1_-vRNA) structures^4,18^, adopting a “closed” conformation (Fig. 2c-d). To investigate how the conformation of B loop affects the relocation of the 3′-RNA template, we created models in which the 3′-RNA template directs towards the catalytic cavity by superimposing the incoming 3′-vRNA template from the transcription initiation state (PDB 5MSG) into the FluPol_H5N1_-cRNA structure, FluPols in intermediate state and transcription pre-initiation state (Fig. 2a-d). These models showed that the B loop with an “open” conformation allows entry of the RNA template into the catalytic cavity (Fig. 2a-b). Conversely, the B loop with a “closed” conformation results in a steric hindrance, preventing entry of the template RNA. These observations were consistent with previous findings that the “closed” conformation of the B loop is incompatible with the presence of a template^28,29^(Fig. 2c-d).

**Fig. 2.**
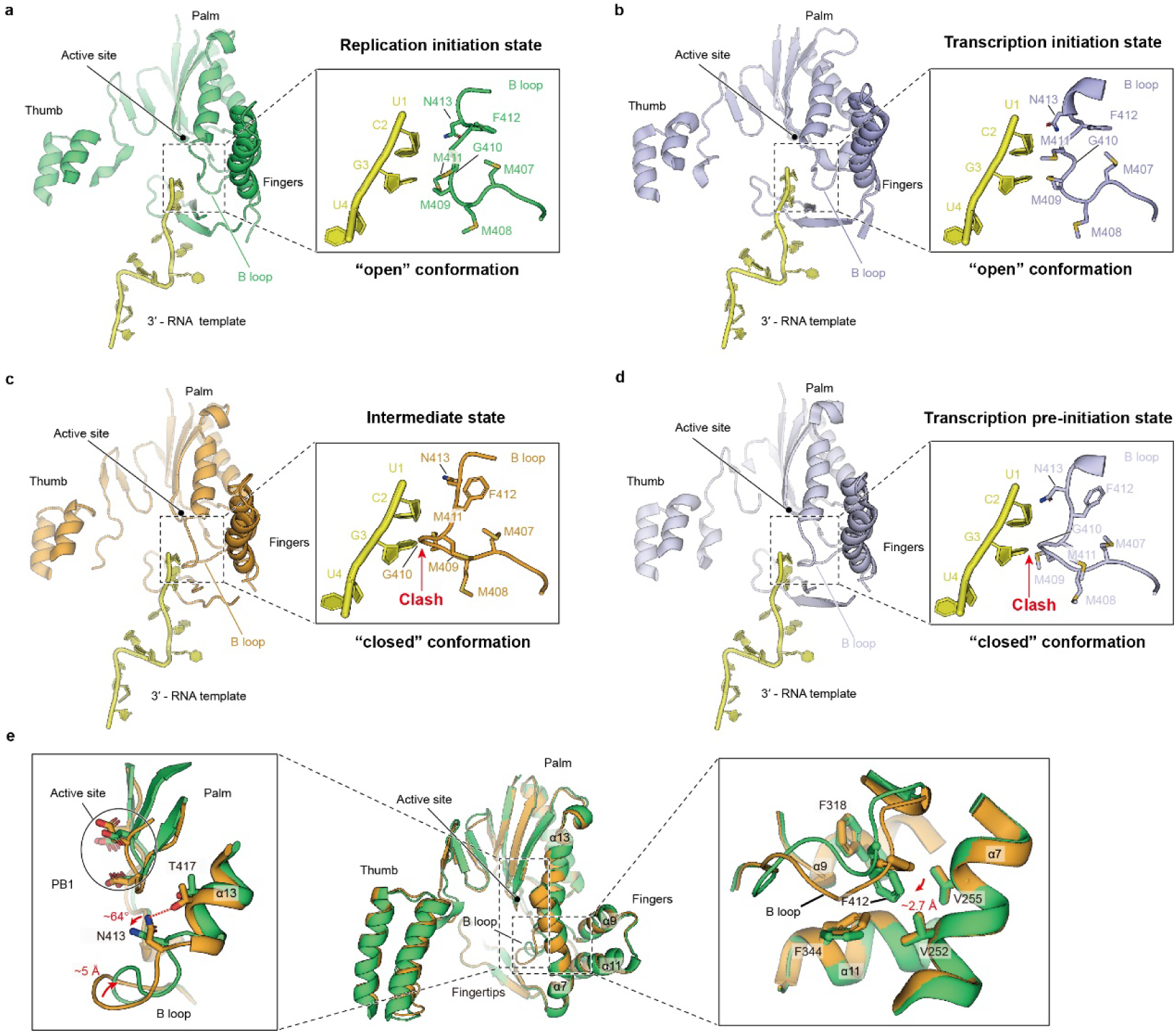
The structural features of the B loop in FluPols. **a-d.** Models of the 3′-RNA template (yellow) directed towards the catalytic cavity in FluPol_H5N1_-cRNA structure (termed the pre-catalytic state, **a**, green), in intermediate state (**c**, orange, PDB 8H69) and transcription pre-initiation state (**d**, blue-white, PDB 4WSB) of FluPols were created by superimposing the RNA template from the transcription initiation state (**b**, light blue, PDB 5MSG), in which the 3′-RNA template has translocated into the catalytic cavity. The B loop adopts an “open” conformation in pre-catalytic state (**a**) and transcription initiation state (**b**), allowing the 3′-RNA template to enter into the catalytic cavity. The B loop adopts a “closed” conformation in the intermediate state (**c**) and transcription pre-initiation state (**d**), causing steric clashes with the 3′-RNA template. **e**. The B loop undergoes a conformational transition from the “closed” (orange) state to the “open” (green) state (**left**). The residue Phe412 is surrounded by a hydrophobic pocket consisting of residues Val252, Val255, Phe318 and Phe344 around the B loop (**right**).

To analyze the conformational transition of the B loop in FluPol_H5N1_ from the “closed” state to the “open” conformation, we compared the B loop in FluPol_H5N1_-cRNA structure with that in the intermediate state structure. The B loop with a “closed” conformation undergoes a retraction of approximately 5 Å against the PB1 finger domain (Fig. 2e, left), with the benzene ring of the residue Phe412 inserting approximately 2.7 Å deeper, fitting into a nearby hydrophobic pocket surrounded by the residues Val252 and Val255 on helix α7, Phe318 on helix α9, and Phe344 on helix α11 of PB1 (Fig. 2e, right). These structural rearrangements of the B loop result in an “open” conformation, enabling entry of the RNA template into the catalytic cavity. Furthermore, we observed that the side chain of the residue Asn413 in the B loop undergoes a flip during this structural transition process. This asparagine, which is conserved in all hand-shaped polymerases, is responsible for recognizing the ribose hydroxyl of incoming NTPs^24^. When the B loop is in the “closed” conformation, the amide of the residue Asn413 side chain forms a hydrogen bond with the hydroxyl group of the residue Thr417 on helix α13 of PB1. By contrast, when the B loop is in the “open” conformation, the side chain of residue Asn413 flips approximately 64° with slight backbone movement towards the catalytic site, positioning it for NTPs recognition (Fig. 2e, left). These findings illustrate that the B loop with an “open” conformation indicates the translocation of the 3′-cRNA template into the catalytic cavity, suggesting the FluPol_H5N1_-cRNA complex represents a state which is ready for replication initiation.

### The Val273 on the PR loop limiting the dynamic the 3′-cRNA for facilitating its realignment

In comparison to all previously resolved structures of FluPol, another feature in the FluPol_H5N1_-cRNA structure is the PR loop (residues 272-276) in the helix-loop-helix motif (residues 269-282) on the PB1 subunit, located above the catalytic site of polymerase (Fig. 3a). Previous report has shown that the PR loop in LACV stabilizes the 3′ terminus of RNA template which enters the catalytic cavity^23^. Given that the PR loop is conserved among polymerases of negative-sense RNA viruses, we speculated that it performs a similar function in influenza polymerase.

**Fig. 3.**
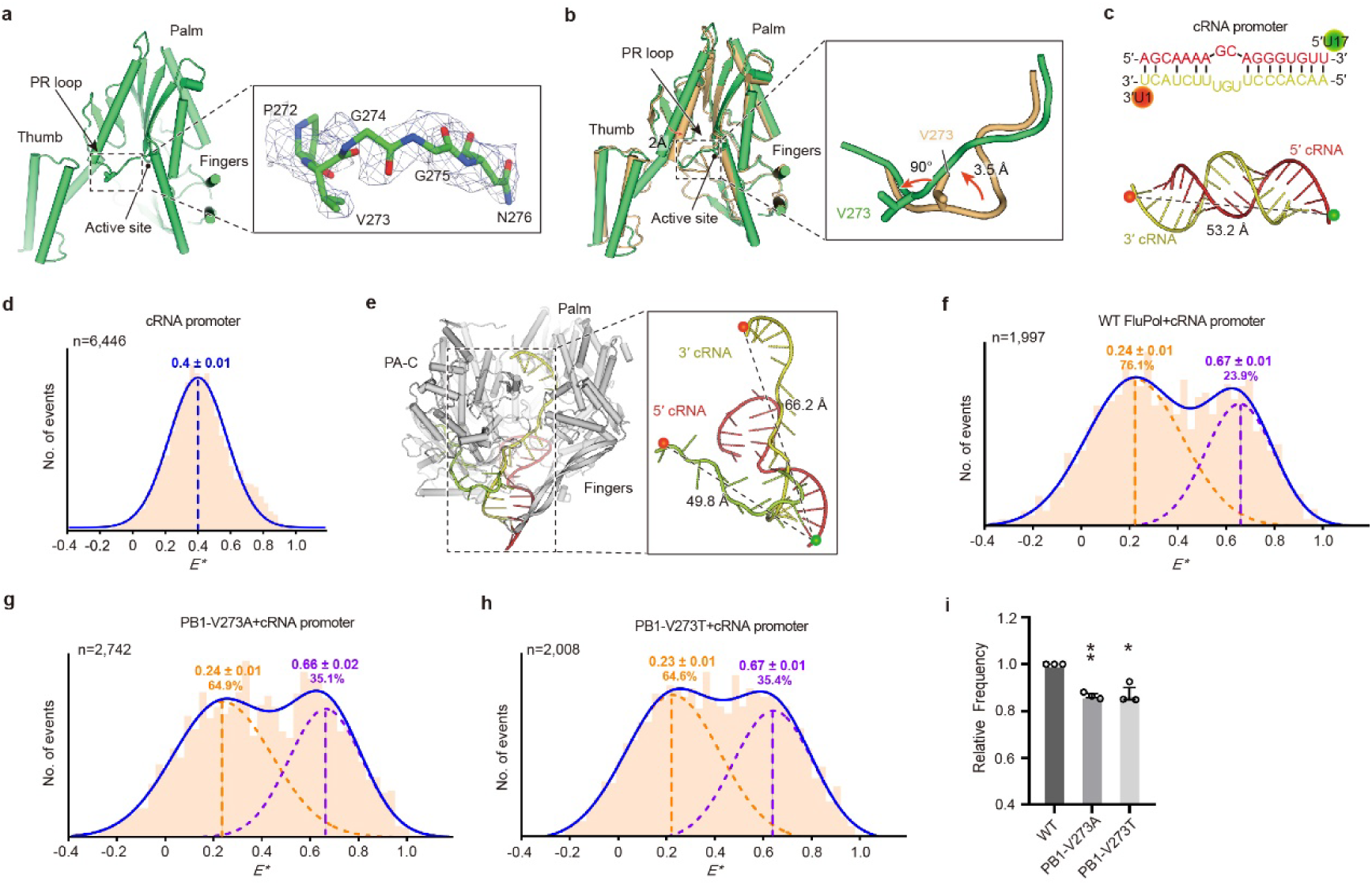
The PR loop plays an important role in stabilization of the 3′-cRNA template. **a.** The location and density map of the PR loop. **b.** Structural comparisons of the PR loop between the FluPol_H5N1_-cRNA (green) and intermediate (orange) structures. **c.** Annealed cRNA promoter labelled with donor and acceptor dyes at positions U17 of 5′-cRNA (17-mer) and U1 of 3′-cRNA (18-mer), respectively (**top**). The model of the fluorophore positions is shown on the double-strand cRNA structure predicted by the Alphafold 3 server. The distances between dyes labelled on the promoters were measured (**bottom**). **d, f-h.** The cRNA promoters labelled with donor and acceptor fluorophores either analyzed alone (**d**), or incubated with FluPol_H5N1_ (**f**), mutant PB1-V273A (**g**) and mutant PB1-V273T (**h**) using single-molecule FRET spectroscopy of diffusing molecules. Efficiency (*E\**) represents the FRET efficiency, *n* represents the number of molecules, and curves that were fitted with Gaussian functions to determine the center of distributions. The ratios of Peak 1 (orange) or Peak 2 (purple) to the total number of FRET events were shown as indicated. **e.** The model of the fluorophore positions was created in PyMOL by superposing the FluPol_H5N1_ 3′-cRNA (17/18-mer) structures predicted by the Alphafold 3 server. These structures exhibit two distinct conformations, one oriented towards the catalytic cavity and the other bound to the second binding site. The distances between dyes labelled on the promoters were measured. **i.** Statistical analysis of the related percentage of molecules from single-molecule FRET data of the cRNA promoters incubated with FluPol_H5N1_. Data are shown as mean ± s.e.m.; *n* = 3 independent experiments. *P* values were calculated by one-way analysis of variance (ANOVA); \**P* < 0.05; \*\**P* < 0.01.

We observed that the PR loop adopts a unique conformation in the FluPol_H5N1_-cRNA structure. In previously reported structures, the PR loop protrudes towards the catalytic site, with the side chain of the residue Val273 adopting an upward conformation away from the catalytic cavity (Fig. S2). In contrast to the intermediate state structure (PDB 8H69), the PR loop in the FluPol_H5N1_-cRNA structure shifts approximately 3.5 Å upward away from the catalytic site, simultaneously pushing an adjacent helix α8 approximately 2 Å away from the cavity, resulting in an approximately 90° flip of the side chain of Val273 towards the catalytic cavity (Fig. 3b). We hypothesized that the residue Val273 in the PR loop, oriented towards the catalytic site, is the key residue for facilitating the 3′ terminus of the RNA template realignment, which may explain why the V273A mutation leads to the generation of failed-realignment products in previous reports^19^.

To validate this hypothesis, we substituted Val273 with alanine (PB1-V273A) or threonine (PB1-V273T), and performed a single-molecule Förster resonance energy transfer (smFRET) method as previously described^30–32^. The results of the smFRET assays showed that the cRNA promoter alone, which folds into a partial complementary double-stranded structure (Fig. 3c), yielded a single FRET distribution (centered at *E\** ∼ 0.4) (Fig. 3d). Upon the addition of FluPols, the 3′ terminus of the cRNA promoter enters the catalytic cavity or binds to the secondary binding site (Fig. 3e), resulting in a bimodal FRET distribution, which is consistent with the observations in previous report^30,31^. When bound to the labelled cRNA promoter, the wild-type FluPol, the PB1- V273A mutant, and the PB1-V273T mutant all produced a low-FRET population (centered at *E\** ∼ 0.24) and a high-FRET population (centered at *E\** ∼ 0.67) (Fig. 3f-h). Owing to the distances between fluorescent dyes being different (Fig. 3e), the low-FRET population corresponded to the conformation of the 3′-cRNA template which directs towards the active site, while the high-FRET population corresponded to that binding in the secondary binding site. Our results showed that the ratios of the low-FRET population to the total number of FRET events for wild-type FluPol, PB1-V273A or PB1-V273T mutants were 76.1%, 64.9% and 64.6%, respectively, while those of the high-FRET population were 23.9%, 35.1% and 35.4%, respectively (Fig. 3f-h). The total number of FRET events in the low-FRET population decreased by approximately 14.8% and 15.2% for the PB1-V273A or PB1-V273T mutants, respectively, compared to the wild-type FluPol (Fig. 3i). These results showed that mutation at Val273 resulted in a decrease in the low-FRET population compared to that of wild-type FluPol, suggesting that the Val273 plays an important role in limiting the dynamic of 3′-cRNA within the catalytic cavity, which is consistent with previous findings^32^. Notably, using the same method, we demonstrated that the Val273 did not restrict the dynamics of 3′- vRNA template (Fig. S3), which is consistent with previous functional studies that the V273A mutant did not affect the replication activity on the vRNA template^19^.

To investigate whether the mutation from valine to alanine could impact the overall conformation of the PR loop and to explore the mechanism by which Val273 limits the dynamic of the 3′-cRNA template, we tested the polymerase activity of the PB1-V273A mutant (which eliminates the side chain) and the PB1-V273T mutant (which is structurally the most similar to alanine) *in vitro*. We expressed the PB1-V273A and PB1-V273T mutants using a Bac-to-Bac expression system (Fig. S4a). To assess whether the PB1-V273A and PB1-V273T mutants affect the transcription activity of FluPol, we performed the *in vitro* cap-dependent transcription assays. Our results showed that the accumulation of transcription products from both the PB1-V273A and PB1-V273T mutants was comparable to that of wild-type FluPol, suggesting that these mutations did not impair the initial step of transcription of FluPol (Fig. 4a). Next, we tested which stage of the replication cycle was affected by these mutations. We assessed whether the PB1-V273A and PB1-V273T mutants affect the replication initiation using *in vitro* dinucleotide synthesis assays as previously described^15^. Our results showed that the ability of both PB1-V273A and PB1-V273T mutants to synthesize terminal or internal pppApG on the vRNA or cRNA template was comparable to that of wild-type FluPol (Fig. 4b). These results indicate that both the PB1-V273A and PB1-V273T mutants do not affect the *de novo* replication initiation of FluPol.

**Fig. 4.**
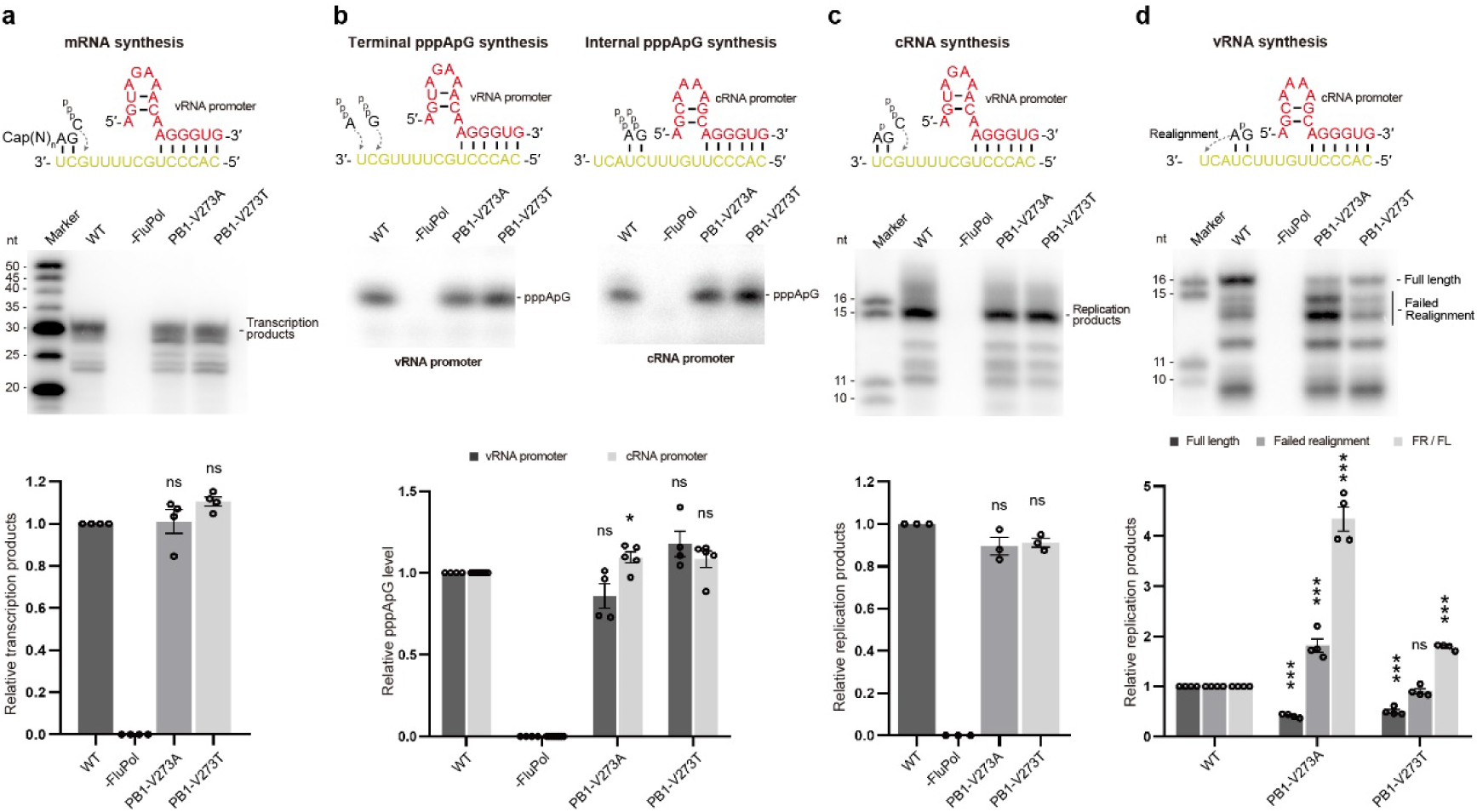
The PR loop is specifically required for the vRNA synthesis *in vitro*. **a**. Effect of the residue Val273 on FluPol_H5N1_ activity in cap-dependent transcription assays. Data are shown as mean ± s.e.m., *n* =4 independent experiments. The *P* values were calculated using one-way ANOVA; ns, no significance. **b.** *In vitro* dinucleotide synthesis assay using the vRNA (**left**) or cRNA (**right**) promoter as a template. Data are shown as mean ± s.e.m., *n* =4 independent experiments. The *P* values were calculated using one-way ANOVA; *, *P* < 0.05, ns, no significance. **c, d** *In vitro* ApG extension assays using the vRNA (**c**) or cRNA (**d**) promoter as a template. Data are shown as mean ± s.e.m., *n* ≥ 3 independent experiments. The *P* values were calculated using one-way ANOVA; ***, *P* < 0.001, ns, no significance.

Furthermore, to investigate whether the PB1-V273A and PB1-V273T mutants affect the replication activity, we performed *in vitro* ApG extension assays using the vRNA or cRNA template. The results showed that the ability of both the PB1-V273A and PB1- V273T mutants to synthesize the cRNA products using the vRNA template was comparable to that of wild-type FluPol (Fig. 4c). However, the ability of the PB1- V273A and PB1-V273T mutants to synthesize the full-length vRNA products was reduced by approximately 63% and 60%, respectively, compared to that of wild-type FluPol. The accumulation of failed-realignment products of the PB1-V273A mutant was increased by 70%, compared to that of wild-type FluPol, while that of the PB1- V273T mutant was comparable to wild-type FluPol. Consequently, the ratios of full-length to failed realignment vRNA products of the PB1-V273A and PB1-V273T mutants significantly increased by 350% and 90%, respectively, compared to that of wild-type FluPol (Fig. 4d). These results indicate that disrupting the hydrophobicity of the residue Val273 impairs the cRNA template realignment process during the vRNA synthesis, suggesting that the residue Val273 limits the dynamic of the 3′-cRNA through hydrophobic effect.

Overall, these findings indicate that the residue Val273 on the PR loop serves to restrict the dynamics of the 3′-cRNA through the hydrophobic effect, thereby facilitating the cRNA template realignment process during internal initiation for vRNA synthesis.

### An asparagine cluster is required for polymerase activity

In addition to the aforementioned distinctions, another notable structural divergence observed in the FluPol_H5N1_-cRNA structure is that the α16, α17 and η7 helices within the thumb domain move approximately 4 Å toward the catalytic cavity and the α8 helix on the palm domain moves approximately 2.5 Å outward the catalytic cavity when compared to the FluPol structures in transcription initiation state (Fig. 5a). Notably, an asparagine cluster, comprising residues Asn504 on the η7 helix, Asn532, Asn533, Asn536 and Asn537 on the α16 helix, as well as Asn276 on the helix-loop-helix motif, was identified above the catalytic cavity (hereinafter referred to as the Asn-cluster) (Fig. 5b). The structural rearrangement of α8, α16 and η7 helices leads to the formation of internal interactions between these asparagine residues. Specifically, the side chain carbonyl group of Asn504 interacts with the main chain amide group of Ser506, while its side chain amide group interacts with the side chain carbonyl group of Asn537 in the FluPol_H5N1_-cRNA structure. The flipping out of Asn276 disrupts the interaction network formed by Asn276, Asn535 (Asn536 in FluPol_H5N1_), Arg144 and Phe218 of the PB2 subunit in transcription state (Fig. 5a).

**Fig. 5.**
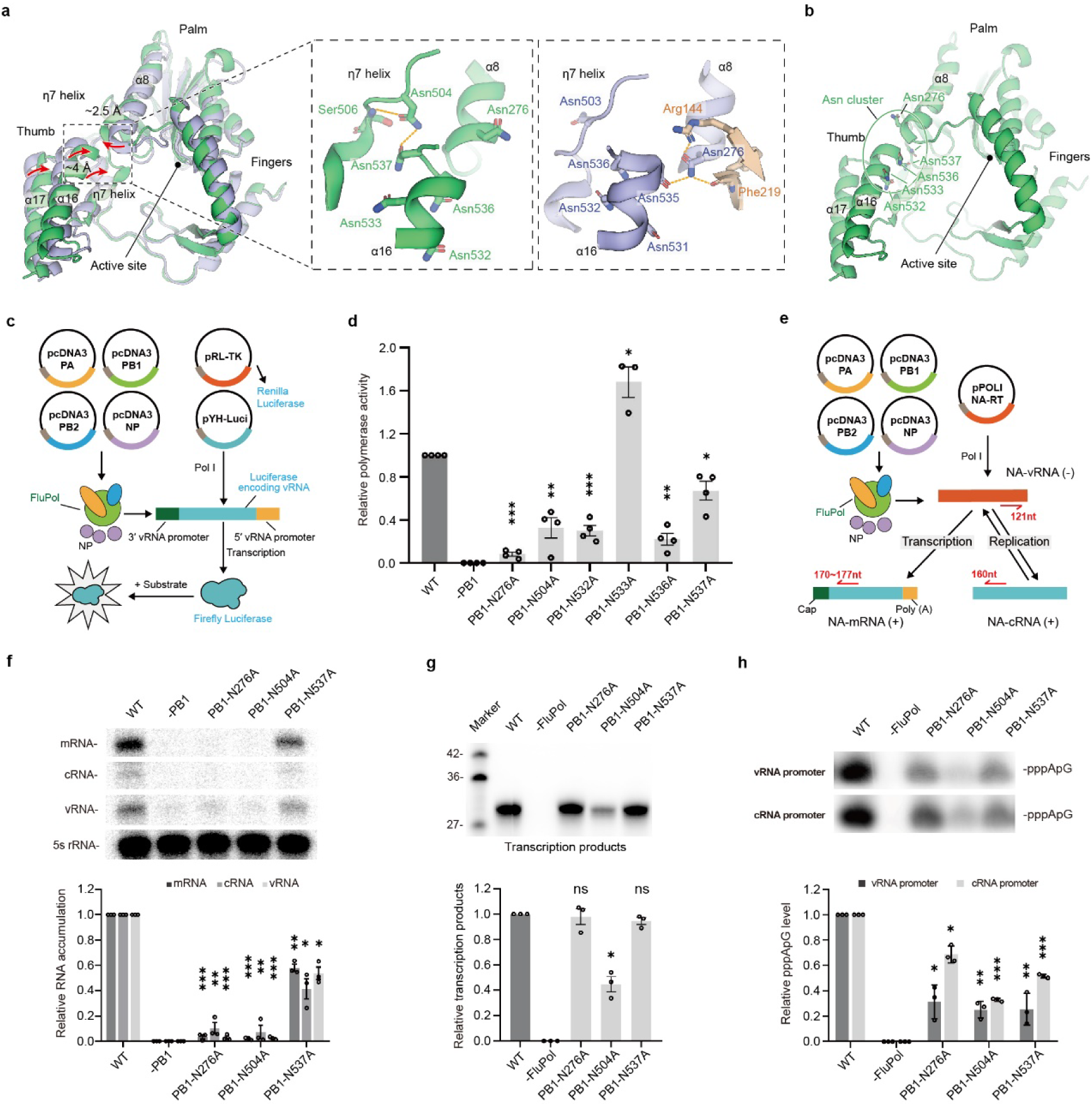
The Asn-cluster is required for polymerase activity of FluPol. **a**. Structural comparisons of the Asn-cluster between the FluPol_H5N1_-cRNA (green) and FluPol in transcription initiation state (PB1 shown in light blue, PB2 shown in orange, PDB 5MSG). Close-up views of the interactions stabilizing the Asn-cluster. Interacting residues are shown in stick representations. **b.** Cartoon representations of the Asn-cluster in the FluPol_H5N1_-cRNA structure. **c, e.** Schemes of polymerase activity assay (**c**) and primer extension assay (**e**). **d.** Effect of mutations at residues Asn276, Asn504, Asn532, Asn533, Asn536 and Asn537 on overall polymerase activity in a minigenome luciferase reporter assay. Data are shown as mean ± s.e.m.; *n* = 4 independent experiments. *P* values were calculated by one-way analysis of variance (ANOVA); \**P* < 0.05, \*\**P* < 0.005, \*\*\**P* < 0.001. **f.** Effect of mutations at the residue Asn276, Asn504 and Asn537 on polymerase activity in a viral RNP reconstitution assay. Data are shown as mean ± s.e.m.; *n* = 3 independent experiments. *P* values were calculated by unpaired two-tailed, one-sample *t*-test; \**P* < 0.05, \*\**P* < 0.005, \*\*\**P* < 0.001. **g.** Cap-dependent transcription assays. Data are shown as mean ± s.e.m.; *n* = 3 independent experiments. *P* values were calculated by one-way ANOVA; \**P* < 0.05, ns, no significance. **h.** *In vitro* dinucleotide synthesis assay. Data are shown as mean ± s.e.m., *n* = 3 independent experiments. The *P* values were calculated using one-way ANOVA; \**P* < 0.05, \*\**P* < 0.005, \*\*\**P* < 0.001.

To test the potential role of the Asn-cluster, we generated point mutations by substituting asparagine with alanine, including the PB1-N276A, PB1-N504A, PB1- N532A, PB1-N533A, PB1-N536A and PB1-N537A mutants. First, we confirmed that these mutants did not affect the expression of FluPol (Fig. S4b). To assess the overall polymerase activities of these mutants, we performed a standard minigenome reporter assay (Fig. 5c). The results showed that all mutants, except for PB1-N533A, significantly decreased the overall polymerase activity compared to that of wild-type FluPol. In contrast, the PB1-N533A mutant exhibited a significant enhancement in polymerase activity (Fig. 5d). These findings indicate that the Asn-cluster is required for polymerase activity.

To investigate whether the internal interactions of the Asn-cluster are required for the transcription or replication processes, we measured the cellular accumulations of mRNA, vRNA, and cRNA using an RNP reconstitution and primer extension assay on the PB1-N276A, PB1-N504A and PB1-N537A mutants, which disrupted the internal interactions of the Asn-cluster (Fig. 5e). Our results demonstrated that the PB1-N276A, PB1-N504A and PB1-N537A mutants all resulted in a significant reduction in the accumulations of mRNA, vRNA, and cRNA, compared to wild-type FluPol (Fig. 5f). These findings suggest that the internal interactions of the Asn-cluster are crucial for transcription or replication processes. However, whether the transcription process and cRNA synthesis were impaired remains unknown, as vRNA serves as the template for transcription and cRNA synthesis.

To further investigate the functions of the internal interactions of the Asn-cluster *in vitro*, we expressed and purified the PB1-N276A, PB1-N504A and PB1-N537A mutants (Fig. S4a). To test whether these mutants affect the transcription activity of FluPol, we performed *in vitro* cap-dependent transcription assays. Our results showed that transcription activity of both the PB1-N276A and PB1-N537A mutants was comparable to that of wild-type FluPol, whereas the transcriptional activity of the PB1- N504A mutant was reduced by approximately 55%. These findings suggest that PB1- N276A and PB1-N537A mutants do not affect the initial step of transcription of FluPol, whereas the PB1-N504A mutant is critical for this process (Fig. 5g).

We then tested whether these mutations affect the replication initiation of FluPol using *in vitro* dinucleotide synthesis assays. Our results showed that the terminal pppApG synthesis ability of all the PB1-N276A, PB1-N504A and PB1-N537A mutants on the vRNA template was reduced by approximately 68%, 75% and 75%, respectively, while that on the cRNA template was reduced by approximately 32%, 77% and 48%, respectively, compared to that of wild-type FluPol (Fig. 5h). These results showed that the PB1-N276A, PB1-N504A and PB1-N537A mutants significantly affected the *de novo* replication initiation of FluPol. These findings indicate that maintaining the internal interactions of the Asn-cluster is crucial for replication initiation of FluPol.

## Discussion

In this study, we report a FluPol_H5N1_-cRNA structure which represents a pre-catalytic state. This structure is characterized by the B loop adopting an “open” conformation, the residue Val273 in the PR loop, and the Asn-cluster protruding towards the catalytic cavity (Fig. S5). These structural features are demonstrated by functional assays that are required for replication initiation of FluPol.

In the FluPol_H5N1_-cRNA structure, we obtained only the core region consisting of PA- C, PB1, and PB2-N subunits, while the highly flexible PA-N and PB2-C domains were absent. This is consistent with previous reports on cRNA-bound FluPol_B_ and cRNA- bound FluPol_D_ structures, which also suggest significant structural variability of FluPol upon cRNA promoter binding^6,10^. Despite the absence of the flexible PA-N and PB2- C, this structure reveals unique conformational characteristics of several key elements within the core region where the replication initiation occurs. The 3′-cRNA, which exhibits high-affinity binding (KD≤1.6 nM) to the secondary binding site on the surface of FluPol^33^, situates at the template entrance channel instead of binding to the secondary binding site in the FluPol_H5N1_-cRNA structure (Fig. 1c). This observation suggests that the 3′-cRNA is oriented toward the catalytic cavity, a feature of FluPol that represents a pre-catalytic state.

In the FluPol_H5N1_-cRNA structure, we observed that the B loop adopts an “open” conformation, which facilitates entry of the RNA template into the catalytic cavity. This conformation is accompanied by a rotation of Asn413 in the B loop towards the catalytic sites, which is critical for NTP recognition during replication initiation. These observations are consistent with previous studies indicating that the B loop plays a critical role in allosteric regulation of polymerase activity by correcting the position of the template nucleotide in the active site and facilitating the binding of the incoming NTP in poliovirus and other viruses^24^.

Although previous studies have shown that the residue Val273 affects the cRNA template realignment process, the specific mechanism remains unknown due to a lack of structural information^19^. In the FluPol_H5N1_-cRNA structure, we observed a unique conformation of the PR loop, characterized by a distinct rotation of residue Val273 towards the active site. This specific conformation was not observed in other resolved FluPol structures. Our smFRET results demonstrate that residue Val273 in the PR loop specifically restricts the dynamic of the 3′-cRNA template but not the vRNA template, as it translocates into the catalytic cavity. These findings are consistent with previous reports indicating restricted dynamics of the first two nucleotides at 3′-cRNA template^32^, suggesting that Val273 in the PR loop, through its conformational change, induces the observed restriction of 3′-cRNA dynamics. Additionally, we observed that the conformation of the residue Val273 in the PR loop of the resolved FluPol_H5N1_-cRNA structure is completely different from that in the cRNA-bound FluPol structure predicted by the AlphaFold 3 server (Fig. S6), indicating that the AI-generated model of FluPol_H5N1_-cRNA may not be entirely accurate, particularly regarding the conformations of critical structural elements.

Furthermore, based on previous results showing that the PB1-V273A mutant affects the cRNA template realignment process^19^, we further revealed that the PB1-V273T mutant, whose hydrophobicity is disrupted, specifically affected vRNA synthesis, resulting in significantly less full-length but more failed-realignment vRNA products, compared to the wild-type FluPol. Notably, mutations on Val273 did not affect their ability to synthesize the dinucleotide pppApG. Based on these results, we propose that Val273 is dispensable for the priming and pppApG synthesis steps of replication initiation. During these steps, both cRNA and vRNA templates utilize analogous mechanisms for pppApG synthesis, with the sole difference being their template positioning due to distinct lengths. For vRNA, the 5′-terminal nucleotides (positions +1 and +2) are positioned in the catalytic site. Thus, directs pppApG synthesis from the template’s very beginning. For cRNA (extend deeper by 2 nucleotides), nucleotides at positions +4 and +5 are positioned in the catalytic sites. Thus, achieves equivalent pppApG synthesis based on the positions +4 and +5. Consistently, the V273A mutation shows no significant change in pppApG synthesis efficiency. Given that the PR loop is strictly conserved across all influenza virus polymerases (Fig. S7) and the mechanism by which the amino acids on the PR loop facilitate the RNA template realignment has been reported in other viruses^23^, it is reasonable to propose that the influenza virus utilizes a similar mechanism.

Moreover, we identified an Asn-cluster above the catalytic cavity in the FluPol_H5N1_- cRNA structure. Our biochemical analyses indicate that the Asn-cluster is required for polymerase activity. Particularly, the residues Asn276, Asn504 and Asn537 are crucial for replication initiation of FluPol. The interactions among Asn504, Asn537, and Ser506 of the thumb domain stabilize the conformation of the η7 helix (loop), which is consistent with previous studies that the η7 loop facilitates replication initiation by stabilizing the priming loop^17,18^. The flipping out Asn276 adopts a conformation beneficial for replication initiation by disrupting the interaction network specifically formed in transcription initiation state. The reorganization of the interactions required for replication initiation indicates that the FluPol_H5N1_-cRNA structure presents a state, conducive to replication initiation. In addition, the mechanistic basis by which the dual role of Asn504 decreases transcription activity remains to be elucidated. Further investigation is needed to elucidate the mechanisms by which residues Asn532 and Asn533 influence polymerase activity, which do not exhibit specific interactions in the FluPol_H5N1_-cRNA structure.

Based on our comprehensive structural and functional analyses, we propose a model for the mechanism of cRNA template realignment during replication initiation of vRNA synthesis from the cRNA template (Fig. 6). As the B loop adopts an “open” conformation, the 3′-cRNA template translocates into the active site. The priming loop, which extends from the thumb domain towards the catalytic sites, provides a stack platform for incoming NTPs incorporation and the B loop facilitates the incoming NTPs binding with the RNA template^4,17,19^. Specifically, the cRNA template is situated near the catalytic site and forms base pairs with the synthesized dinucleotide pppApG at positions +4 and +5. It was reported that the FluPol in pre-catalytic state binds to an apo polymerase and forms a dimer^19^. Dimerization triggers the downward movement of the helix buddle composed of PB1-C and PB2-N, causing the priming loop to move downward and thereby driving the cRNA template to backtrack. Since the first two nucleotides of 3′-cRNA template inserts into a deeper location within the catalytic cavity. We propose that the Val273 might limit the dynamic of the first two overhanging nucleotides of the 3′- cRNA by flipping toward the active site, thereby directly promoting the cRNA template backtrack or indirectly facilitating the priming loop to drive the template backward more efficiently. Consequently, the synthesized pppApG realigns to the cRNA template, forming base pairs at positions +1U and +2C^10,19^. Simultaneously, the template-product base pairs progressively elongate to synthesize complete genomic RNA upon NTPs incorporation.

**Fig. 6.**
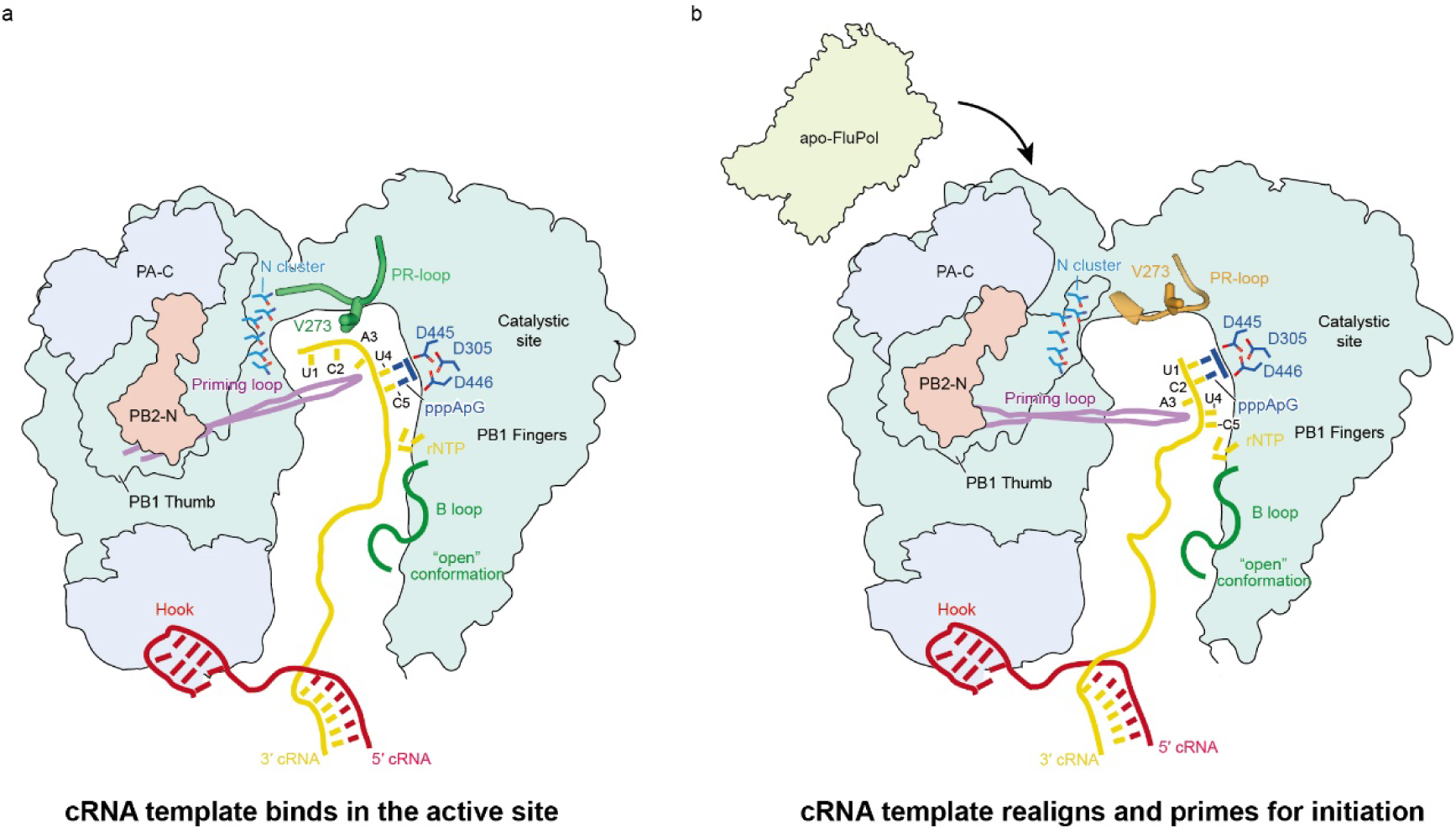
Model of the residue Val273 of the PR loop in template realignment during replication initiation on a cRNA template. **a.** The B loop of FluPol in an “open” conformation allows the entry of the 3′ terminus of the cRNA template. The 3′-cRNA template is stabilized by the residue Val273 in the PR loop at a deeper location of the catalytic cavity and facilitates its priming and realignment for initiating vRNA synthesis. **b.** The dimerization of the replicase and an apo FluPol triggers the conformational changes of the priming loop, resulting in backtracking of the cRNA template.

In summary, our results provide a model for replication initiation during the vRNA synthesis by FluPol, which not only offers further insights into the mechanisms of replication internal initiation, but also holds potential value for anti-influenza drug development.

## Acknowledgements

We thank staff in the Center of Cryo-Electron Microscopy, Zhejiang University for their assistance during data collection. We thank Dr. Gang Ji, Dr. Xiaojun Huang, Dr. Boling Zhu, Mr. Longlong Zhang, Mr. Deyin Fan and Mr. Bingxuan Huangfu at Center for Biological Imaging (CBI), Institute of Biophysics, Chinese Academy of Science for their assistance during data collection and Ms. Tongxin Niu for her computational assistance. We thank the use of Cryo-EM instruments in the Cryo-EM Facility Center of Southern University of Science and Technology. We thank Professor Xiaofeng Zheng (School of Life Sciences, Peking University) for the instruction of the viral functional analysis experiment. We thank Professor George Brownlee’s lab for providing us the pcDNA-PA, pcDNA-PB1, pcDNA-PB2, pcDNA-NP and pPOLI-NA plasmids. We thank Dr. HongJie Zhang (Core Facility for Protein Research, Institute of Biophysics, Chinese Academy of Sciences), Dr. Chengguo Yao and Guangyun Lin (Zhongshan School of Medicine, Sun Yat-Sen University) for assisting us with autoradiography experiments. We thank Mr. Wolun Zhang and Mr. Dunjin Zheng (LightEdge Technologies LTD) for assisting us with the sm-FRET experiments. The work was supported by the grants from Shenzhen Science and technology planning project (project No. JCYJ20220818102017035, ZDSYS20220606100803007 to Y.L.); ’Pearl River Talents Plan’ Innovation and Entrepreneurship Team Project of Guangdong Province (project No.2019ZT08Y464 to Y.L.), National Key R&D Program of China (project No. 2022YFE0210000 to Y.L., 2023YFC2606400 to H.L.); National Natural Science Foundation of China (project No.82071346 to Y.L., 32271321 to Z.W., 82373883 to H.L.); Natural Science Foundation of Guangdong Province (project No.2023A1515010245 to Z.W.); Young Faculty Development Program of Sun Yat-sen University (project No.24qnpy090 to Z.W.) and Jointinnovation project of Guangdong-Hong KongMacao, Grant/Award Number: 2022A0505020023.

## Contributions

Y. W., M. L. T. S., Y. B., S. H. performed the experiment. Y. W., M. L., H. Li., Y. Liu., and H. Liang. wrote the manuscript. All authors reviewed the results and approved the final version of the manuscript.

## Materials and Methods

### Cells and Plasmids

Human embryonic kidney 293T (HEK-293T) cells and Madin-Darby canine kidney (MDCK) cells were cultured in Dulbecco’s Modified Eagle’s Medium (DMEM) and Minimum Essential Medium (MEM), respectively. Both media were supplemented with 10% fetal bovine serum and 1% penicillin-streptomycin. Cells were grown at 37 °C in a 5% CO_2_ atmosphere. Plasmids pcDNA-PA, pcDNA-PB1, pcDNA-PB2 and pcDNA-NP expressing influenza A/WSN/33 virus proteins, as well as the vRNA- expressing plasmids pPOLI-NA, were kind gifts from Professor George Brownlee’s lab. The codon-optimized sequences for three subunits of avian influenza A/Goose/Guangdong//3/1997 (H5N1) virus polymerases were synthesized (GenScript) and cloned into pFastBac expression plasmid for polymerase expression.

### Protein purification

The A/Goose/Guangdong/3/1997 (H5N1) influenza polymerase complex was expressed in the insect cell line High Five after being infected by PA-, PB1-, and PB2- expressing viruses for 60h. Insect cells were collected and resuspended in lysis buffer (50 mM HEPES pH 7.8, 500 mM NaCl, 20 mM imidazole, 10% (*v/v*) glycerol) supplemented with 0.5 mM phenylmethanesulfonyl-fluoride (PMSF), and lysed using a high-pressure homogenizer. The lysates were centrifuged at 16,000 × *g* for 20 min at 4℃ to remove cell debris. The supernatant was incubated with Ni-NTA beads at 4℃ for 2 h. The beads were collected, washed five times and eluted with buffer containing 40 mM HEPES (pH 7.8), 300 mM NaCl, 400 mM imidazole and 5% (*v/v*) glycerol. The eluted proteins were concentrated and purified by heparin affinity chromatography (Cytiva) using buffer A containing 30 mM HEPES (pH 7.8), 400 mM NaCl, 1 mM dithiothreitol and 5% (*v/v*) glycerol and buffer B containing 30 mM HEPES (pH 7.8), 1.2 M NaCl, 1 mM dithiothreitol and 5% (*v/v*) glycerol. The fractions were further subjected to size-exclusion chromatography (Superdex-200, Cytiva) in buffer C containing 30 mM HEPES (pH 7.8), 300 mM NaCl, 1 mM dithiothreitol and 5% (*v/v*) glycerol. The fractions containing the FluPol complex were verified by SDS-PAGE and stored in aliquots at -80℃ for further use.

### Complex preparation

The FluPol_H5N1_-cRNA complex was prepared as previously described^18^. In brief, the 45nt cRNA template whose termini contain conserved 3′ and 5′ promoters (45-mer: AGCAAAAGCAGGGUGUUAAAGUUCAAAAACACCCUUGUUUCUACU) were produced *in vitro*^34^. The purified FluPol_H5N1_ was incubated with cRNA template synthesized *in vitro* at a molar ratio of 1:1.2 on ice for 30 minutes. The cRNA-bound FluPol_H5N1_ monomer with a molecular mass of approximately 250 kD was separated from the mixture on a Superdex 200 column in a buffer containing 20 mM HEPES (pH 7.8), and 300 mM NaCl. Peak fractions were then collected and ultracentrifuged at 15,000 × *g* for 30 minutes at 4°C immediately prior to freezing grids.

### Cryo-EM data acquisition and image processing

The Quantifoil R1.2/1.3 Au 300-mesh holey carbon grids (Quantifoil, Großlöbichau Germany) was made hydrophilic by glow discharging for 60 s in a Gatan Solarus 950. Four microliters of freshly purified FluPol_H5N1_-cRNA complex (0.1 mg/mL) were loaded into grids for preparation of cryo grids using a Vitrobot Mark IV (Thermo Fisher Scientific) at 4°C with a blotting time of 4s (force 2, humidity 100%).

The Cryo-EM of FluPol_H5N1_-cRNA complex were collected on a 300 kV Titan Krios (Thermo Fisher Scientific) electron microscopy equipped with a Gatan K2 Summit direct electron detector (Gatan). Movie stacks were automatically collected using the SerialEM software^35^ with the beam-image shift method at a calibrated magnification of 29,000×, corresponding to a pixel size of 1.014 Å/pixel. Each micrograph stack was exposed for 5.4 s with a total of 32 frames with a preset defocus range from -1.5 μm to -3.5 μm in counting mode. The total dose rate per stack was approximately 50 e/Å^2^. The image processing workflow is depicted in Figure S1. A total of 4,299 micrographs were collected in this session.

The micrographs were first processed separately, including movie alignment using RELION’s motion correction algorithm based on MotionCor2^36^, CTF estimation using Gctf software^37^ and particle picking using auto-picking mode in RELION^38^. A total of 1,591,950 particles were automatically picked and processed with multiple rounds of reference-free 2D classification in RELION. Approximately 781,355 particles were selected and further subjected to global angular searching 3D classification. After several rounds of 3D classifications, a total of 144,735 particles with well-defined structural features were selected and further submitted into RELION_REFINE program which gives rise to a 3.37 Å cryo-EM map. Applying a soft-edge mask and *B* factor sharpening generated a final 2.98 Å cryo-EM map based on the gold-standard Fourier shell correlation 0.143 criterion. Local resolution of the cryo-EM maps of FluPol_H5N1_ complexes was evaluated using ResMap^39^ in RELION.

### Model building

The initial model of FluPol_H5N1_-cRNA was built using the structures of FluPol_A_ bound to vRNA promoters as references (PDB IDs 8H69 and 6RR7). The core region of FluPol was first fitted into the cryo-EM map as a rigid body using UCSF Chimera^40^. Most regions were fitted well into the density; however, we noticed that some regions, like PB2 N-terminal and PB1 thumb, fitted poorly because these regions have undergone a large rotation comparing to those in the vRNA-bound structures. We thus separately fitted these regions using Chimera. In the FluPol_H5N1_-cRNA map, the PB2 N-terminal (residues 61-82, 90-103) and part of PB1 thumb (residues 617-631, 661- 668) have a lower local resolution. This only allowed to trace the main-chain backbone, however, the side chains in these regions were ambiguously built. In the PB2 subunit, we observed two segments of density, corresponding to the two helices, residues 43-52 and 110-124, in the PB2 N-terminal. However, these two local densities were unmodeled due to the map discontinuity. In the PB1 subunit, the PR loop containing several Gly residues and V273 has only few side-chain features and was more flexible than other rigid parts around the active site. The map supported model building of the main-chain backbone of the PR loop and the side-chain of V273. The model was further refined using PHENIX.REAL_SPACE_REFINE program^41^ in PHENIX with chemical restraints including secondary structure and geometry restraints and checked, rebuilt manually in Coot^42^. Model geometry and clash scores were checked using MolProbity^43^.

### Minigenome reporter assay

The polymerase activity can be estimated by a dual-luciferase reporter gene assay as previously described ^44,45^. HEK293T cells were co-transfected with plasmids encoding the wild-type or mutant PB1 (50 ng), PB2 (50 ng), PA (50 ng), and NP (50 ng) proteins, alongside a *firefly* luciferase reporter plasmid pYH-Luci (50 ng) and an internal control plasmid pRL_TK for *Renilla* luciferase (10 ng). After 24 hours of transfection, luciferase activity was measured using a Dual-Glo™ Luciferase Assay System (Promega). *Firefly* and *renilla* luciferase bioluminescence were measured using a Synergy HT plate reader (BioTek, VT, USA). *Firefly* luciferase expression was normalized to *Renilla* luciferase expression (relative luciferase activity). The activity of the wild-type polymerase was set as the baseline at 100%, and the activities of the mutant variants were determined relative to this baseline. Data were analyzed using Prism 8 (GraphPad).

### Oligonucleotide radiolabeling

For the primer extension assay, primers were radiolabeled with [γ-^32^P] ATP. 1 μL of oligonucleotide at 10 μM was mixed with 2 μL [γ-^32^P] ATP (3,000 Ci/mmol, 10 mCi/ml, Perkin Elmer), 1 μL T4 polynucleotide kinase enzyme (NEB), and 1 μL T4 polynucleotide kinase buffer in a total reaction volume of 10 μL. The labelled oligonucleotides were purified using a Qiaquick nucleotide removal kit (QIAGEN), following the manufacturer’s instructions. The purified oligonucleotides were eluted in 30 μL of nuclease-free water (TaKaRa) and stored at -20°C for future use.

### RNP reconstitution and primer extension analysis

Approximately 1×10^6^ HEK-293T cells in 6-well plates were transfected with 1 μg each of pcDNA-PA, pcDNA-PB1, pcDNA-PB2, pcDNA-NP and pPOLI-NA-RT plasmid encoding neuraminidase vRNA segment using PEI (polyscience) according to the manufacturer’s instructions. Cells were harvested 36 hours after transfection. Total RNA was extracted using TRI Reagent (Sigma-Aldrich) according to the manufacturer’s instructions. The accumulation of viral mRNA, cRNA and vRNA was analyzed by primer extension assay. Briefly, RNA was reverse transcribed using the SuperScript III reverse transcriptase (Invitrogen) and ^32^P-labeled primers against positive sense mRNA, cRNA primers (detected the 5′-end portions of viral mRNAs and cRNAs), negative sense vRNA primer and a 5S rRNA primer against cellular 5S rRNA included as a loading control. Products were analyzed by 8%, 8 M Urea-PAGE and visualized by a phosphor-imaging scanner (Typhoon FLA7000IP). Viral RNA signals were quantified using Image J and normalized to the 5S rRNA loading control. Input vRNA signal, estimated from the “-PB1” control, was subtracted from all subsequent lanes. Data were analyzed using Prism 8 (GraphPad)

### *In vitro* dinucleotide synthesis assays

To test the ability of FluPol to synthesize the dinucleotide, 0.4 µM purified recombinant polymerase was incubated with either 0.5 μM 5′-vRNA (16-mer) and 3′-vRNA (15-mer) or 5′-cRNA (15-mer) and 3′-cRNA (16-mer) in the presence of 5 mM MgCl_2_, 1 mM DTT, 2 U µL^-1^ RNasin (Promega), 1 mM ATP, 0.1 µL [α-^32^P] GTP (3,000 Ci mmol^-1^; Perkin-Elmer) and incubated at 30°C for 3 h (RNA sequences shown in table S1). The reactions were quenched at 95°C for 10 min and mixed with an equal volume of loading buffer (80% formamide, 1 mM EDTA, 0.1% bromophenol blue and 0.1% xylene cyan). Samples were then heated again at 95°C for 5 min and cooled immediately on ice-water mixture for 3 min. Finally, the samples were analyzed by a 20%, 8 M Urea-PAGE and visualized by a phosphor-imaging scanner (Typhoon FLA7000IP). All data were analyzed using Image J and Prism 8 (GraphPad).

### *In vitro* ApG extension assays

To test the ability of FluPol to extend from an ApG dinucleotide, 0.4 µM purified recombinant polymerase was incubated with either 0.5 μM 5′-vRNA (16-mer) and 3′- vRNA (15-mer) or 5′-cRNA (15-mer) and 3′-cRNA (16-mer) in the presence of 5 mM MgCl_2_, 1 mM DTT, 2 U µL^-1^ RNasin (Promega), 0.5 mM ApG, 1 mM ATP/UTP/CTP, 1 µM GTP, 0.1 µL [α-^32^P] GTP (3,000 Ci mmol^-1^; Perkin-Elmer) and incubated at 30°C for 3 h (RNA sequences shown in table S1). The reactions were quenched at 95°C for 10 min and mixed with an equal volume of loading buffer (80% formamide, 1 mM EDTA, 0.1% bromophenol blue and 0.1% xylene cyan). Samples were then heated again at 95°C for 5 min and cooled immediately on ice-water mixture for 3 min. Finally, the samples were analyzed by a 20%, 8 M Urea-PAGE and visualized by a phosphor- imaging scanner (Typhoon FLA7000IP). All data were analyzed using Image J and Prism 8 (GraphPad).

### *In vitro* cap-dependent transcription assays

To test the transcription ability of FluPol, 0.4 µM purified recombinant polymerase was incubated with 0.5 μM 5′-vRNA (16-mer) and 3′-vRNA (15-mer) in the presence of 5 mM MgCl_2_, 1 mM DTT, 2 U µL^-1^ RNasin, 2.5 µM capped 16-nucleotide RNAs (5ʹ- m^7^GpppGAAUGCUAUAAUAGC-3ʹ, Trilink), 1 mM ATP/UTP/CTP, 1 µM GTP, 0.1 µL [α-^32^P] GTP (3,000 Ci mmol^-1^, Perkin-Elmer) at 30°C for 3 h (RNA sequences shown in table S1). The reactions were quenched at 95°C for 10 min and mixed with an equal volume of loading buffer (80% formamide, 1 mM EDTA, 0.1% bromophenol blue and 0.1% xylene cyan). Samples were then heated again at 95°C for 5 min and cooled immediately on ice-water mixture for 3 min. Finally, the samples were analyzed by a 20%, 8 M Urea-PAGE and visualized by a phosphor-imaging scanner (Typhoon FLA7000IP). All data were analyzed using Image J and Prism 8 (GraphPad).

### Single-molecule Förster resonance energy transfer (smFRET) experiment

The single-molecule FRET experiments were performed as previously described^30–32,46^. The 5ʹ-vRNA (18-mer) or the 5ʹ-cRNA (17-mer) was labelled with cy3 (donor) at U18 or U17, respectively, while the 3ʹ-vRNA (17-mer) and the 3ʹ-cRNA (18-mer) were both labelled with cy5 (acceptor) at U1 of the 3′ end. These labelled RNAs were synthesized commercially (BGI, Tabel S1). For the samples of RNAs alone, equimolar amounts of the 5ʹ-cRNA (vRNA) and the 3ʹ-cRNA (vRNA) RNA were mixed at a final concentration of 100 nM for 5 min at 95℃ in buffer A (50 mM HEPES (pH8.0), 500 mM NaCl, 10 mM MgCl_2_, 1mM DTT and 5% glycerol) and cooled immediately on ice-water mixture for 3 min, followed by diluted into buffer B (50 mM HEPES (pH8.0), 100 mM KCl, 10 mM MgCl_2_, 1 mM DTT and 5% glycerol) to a final RNA concentration of 150 pM for confocal analysis. For the samples of RNAs bound to FluPols, the Fluorescently labelled RNAs at a final concentration of 1 nM were incubated with FluPols at a final concentration of 100 nM for 15 min at 30℃ in buffer A, followed by diluted into buffer B to a final RNA concentration of 150 pM for confocal analysis.

The smFRET experiments were performed using a commercial bench-top fluorescence correlation spectrometer (CorTectorTM SX200, LightEdge Technologies Limited, China) equipped with a 60× 1.2NA UPLANSAPO water immersion objective (Olympus, Japan) and two (532nm, 638 nm) continuous-wave solid-state lasers (Pavilion Integration Corporation, China). The instrument is based on a confocal optical pathway, using a multimode optical fiber with an inner core diameter of 50 μm as the confocal pinhole. The 532 nm laser or 638 nm laser was used to excite the dyes, and emission light in the spectral range 555-615 nm or 650-900 nm was detected by an optical fiber-coupled single-photon counting avalanche photodiode detector (SPCM- 800-14-FC, Excelitas Technologies Corporation, USA). Based on their photon arrival time, fluorescence photons were classified as originating from donor-based or acceptor-based excitation.

The smFRET data were analyzed by in-house scripts based on the FRETBursts software^47^. Bursts were extracted from three photon streams using “sliding windows” with a window size of *m* = 10, ensuring that the count rate was *F* = 6 times higher than the background level. Additionally, bursts with a total photon count of less than 30 and those containing the were excluded from the analysis. Each fluorescent burst exceeding a specified threshold allows calculation of the apparent FRET efficiency (*E\**) and the fluorophore stoichiometry (*S*), resulting in a two-dimensional histogram. While *E\** indicates the donor-acceptor species, *S* distinguishes molecular species by their relative labeling ratio of green to red fluorophores. A low S of < 0.2 indicates the acceptor-only populations, while a high *S* of > 0.8 corresponds to donor-only populations. One- dimensional *E\** distributions for donor–acceptor species were obtained by using a 0.2 < *S* < 0.8 threshold. The *E\** distributions were fitted using a Gaussian function, thereby determining the mean *E\** value for a given distribution along with the corresponding standard deviation. The data were corrected the *γ* factors (*γ* = 1, a detection correction factor) prior to Gaussian fitting of the *E\** distributions.

### Western blotting analysis

Total protein extracts were clarified by centrifugation at 13,000 × *g* for 15 min at 4°C, and supernatants were mixed with SDS sample buffer and boiled for 10 min. Equal volumes of total proteins were separated by 10% SDS-PAGE for western blotting. Rabbit polyclonal anti-PB1 antibody (both at 1:3,000 dilution, Invitrogen) was used to blot PB1 protein. A β-actin mouse monoclonal antibody (1:1,000 dilution, CST) was used as the internal control. Goat anti-rabbit and goat anti-mouse antibodies conjugated to horseradish peroxidase (HRP) were used as secondary antibodies (1:3,000 dilution, CST). Bands were visualized by chemiluminescence using SuperSignal West Pico Chemiluminescent substrate (Thermo) on a Tanon 5200 Imaging System (Tanon).

### Statistics summary

All experiments were repeated at least three times independently. Statistical analyses were performed using GraphPad Prism 8. Statistical data are presented as mean ± s.e.m. *P* values were calculated by unpaired, two-tailed, one-sample *t* test or one-way ANOVA.

## Reporting summary

Further information on research design is available in the Nature Portfolio Reporting Summary linked to this article.

## Data availability

The cryo-EM maps of FluPol_H5N1_-cRNA complexes have been deposited in the Electron Microscopy Data Bank under accession number EMD-60634. The coordinate for the atomic model of FluPol_H5N1_-cRNA complex has been deposited in the Protein Data Bank under code number 9LCK.

