## Supplementary information for "Cryo-EM structure of influenza polymerase bound to the cRNA promoter provides insights into the mechanism of virus replication"

### Supplementary Figure and Table

### Figure S1

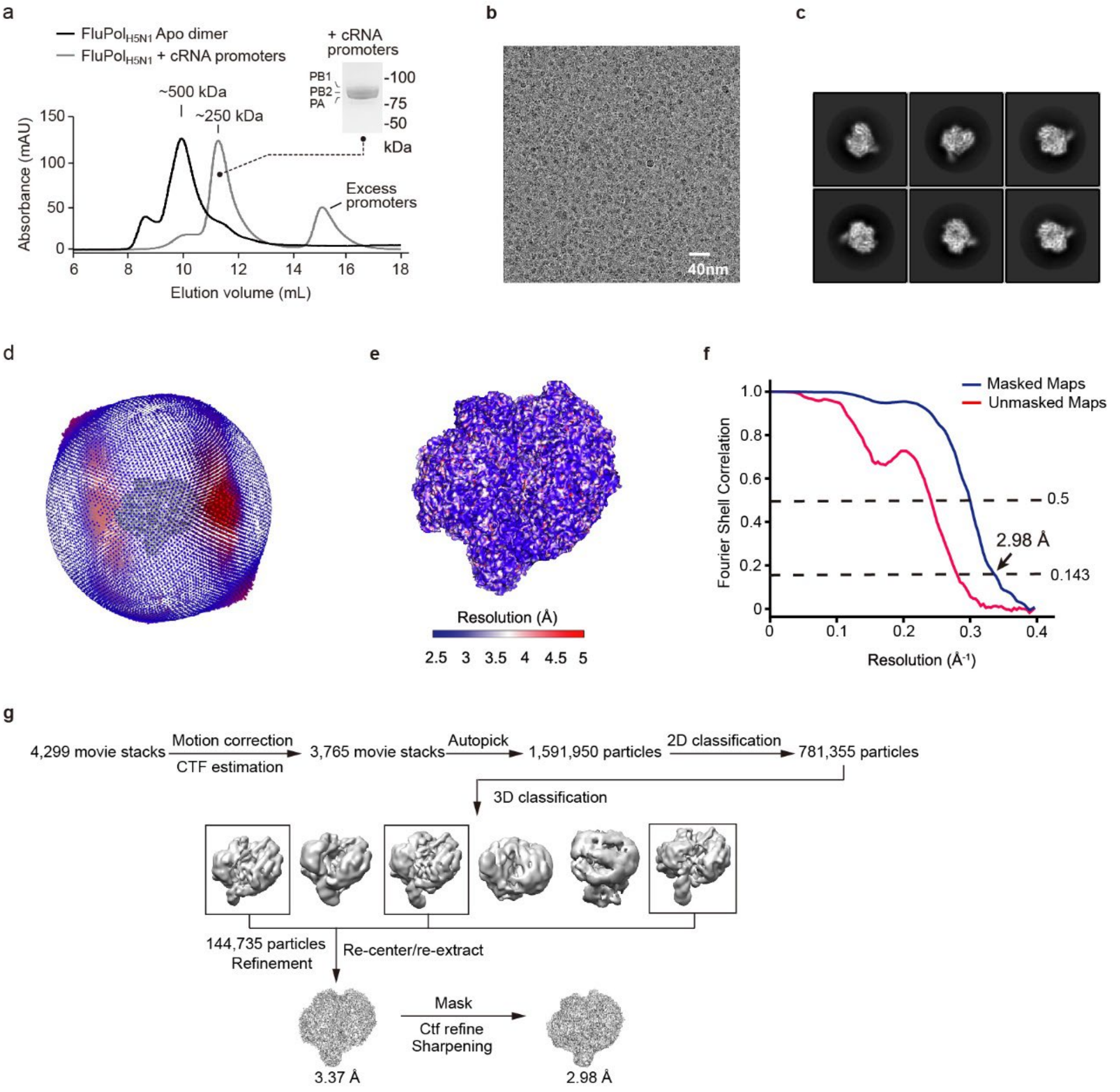


**Figure. S1 Cryo-EM analyses of FluPol$_{H5N1}$-cRNA structure. a.** Size-exclusion chromatography and SDS–PAGE profiles of FluPol$_{H5N1}$ in the apo state (shown in black line) and cRNA promoters-bound state (shown in gray line). **b**. A representative cryo-EM micrograph of FluPol$_{H5N1}$-cRNA structure. **c**. Representative 2D average image of FluPol$_{H5N1}$-cRNA complex particles. **d**. Euler angle distributions of FluPol$_{H5N1}$-cRNA complex in the final 3D reconstruction. **e**. Local resolution evaluations of cryo-EM maps of FluPol$_{H5N1}$ complex by ResMap. **f.** The gold-standard Fourier shell correlation (FSC) curves were calculated at 0.143 and 0.5 cut-off values. **g**. Image processing processes for the 3D reconstruction of FluPol$_{H5N1}$-cRNA structure.

### Figure S2

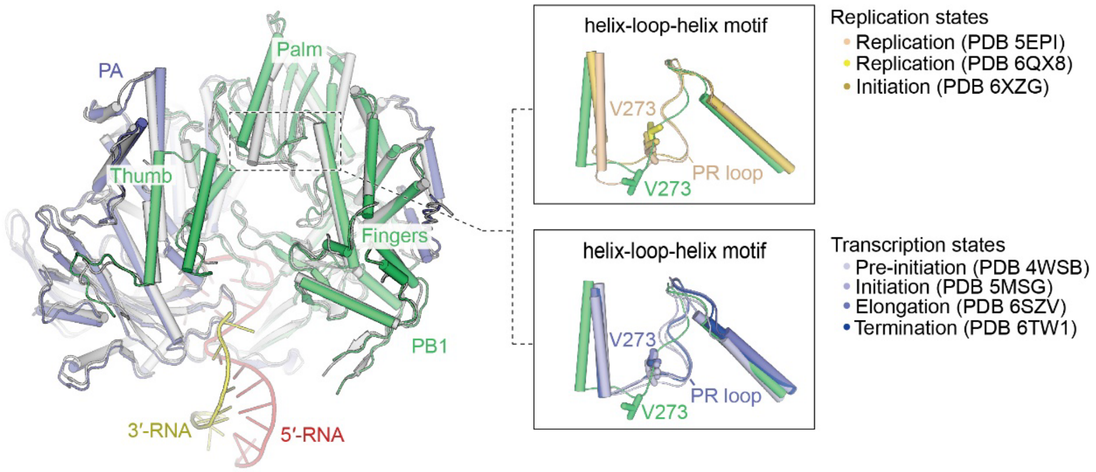


**Fig. S2 The conformation of the helix-loop-helix motif on the PB1 subunit in different states of FluPols.** The residue Val273 flips towards the catalytic cavity in the $FluPol_{H5N1}$-cRNA structure, but adopts an upward conformation away from the catalytic cavity of FluPols in replication initiation state (yellow orange, PDB 6XZG), replication states (yellow, PDB 6QX8; wheat, PDB 5EPI), transcription pre-initiation state (blue white, PDB 4WSB), transcription initiation state (light blue, PDB 5MSG), transcription elongation state (slate, PDB 6SZV) and transcription termination state (blue, PDB 6TW1).

**Figure S3**

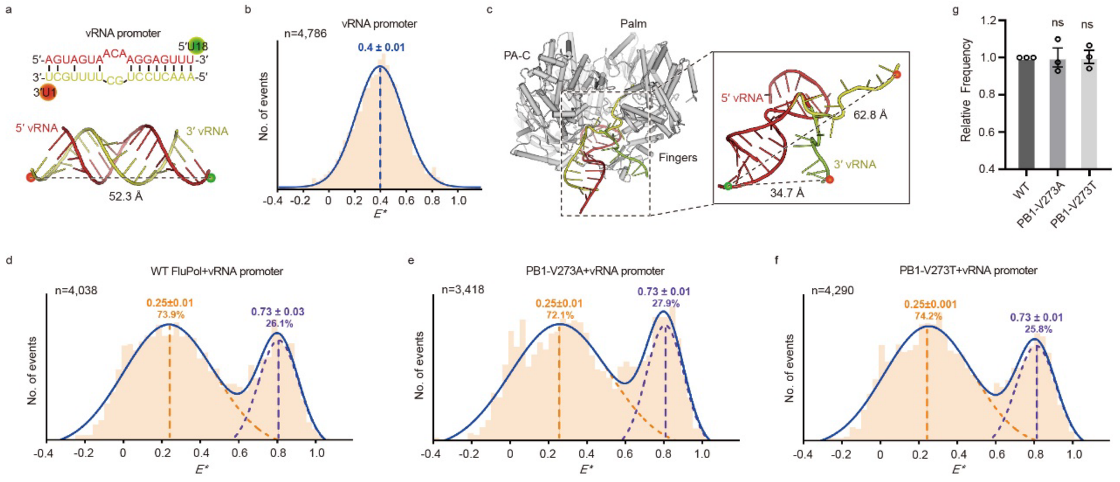


**Fig. S3 The effect of Val273 on the stability of the 3′-vRNA template. a.** Annealed vRNA promoters labelled with donor and acceptor dyes at positions U18 of 5′-vRNA (18-mer) and U1 of 3′-vRNA (17-mer), respectively (**top**). The model of the fluorophore positions is shown on the double-strand vRNA structure predicted by the Alphafold 3 server. The distances between dyes labelled on the promoters were measured (**bottom**). **b, d-f.** Annealed vRNA promoters labelled with donor and acceptor fluorophores either analyzed alone (**b**), or incubated with FluPol$_{H5N1}$ (**d**), PB1-V273A mutant (**e**) or PB1-V273T mutant (**f**) using single-molecule FRET spectroscopy of diffusing molecules. Efficiency ($E$*) represents the FRET efficiency, $n$ represents the number of molecules and curves that were fitted with Gaussian functions to determine the center of distributions. The ratios of Peak 1 (orange) or Peak 2 (purple) to the total number of FRET events are shown as indicated. **c.** The model of the fluorophore positions was created in PyMOL by superposing the FluPol$_{H5N1}$ 3′-vRNA (18/17-mer) structures predicted by the Alphafold 3 server. These structures exhibit two distinct conformations, one oriented towards the catalytic cavity and the other bound to the surface of the FluPol. The distances between dyes labelled on the promoters were measured. **g.** Statistical analysis of the related percentage of the molecules from single-molecule FRET data of the vRNA promoters incubated with FluPol$_{H5N1}$. Data are shown as mean ± s.e.m.; $n$ = 3 independent experiments. $P$ values were calculated by one-way analysis of variance (ANOVA); ns, no significance.

**Figure S4**

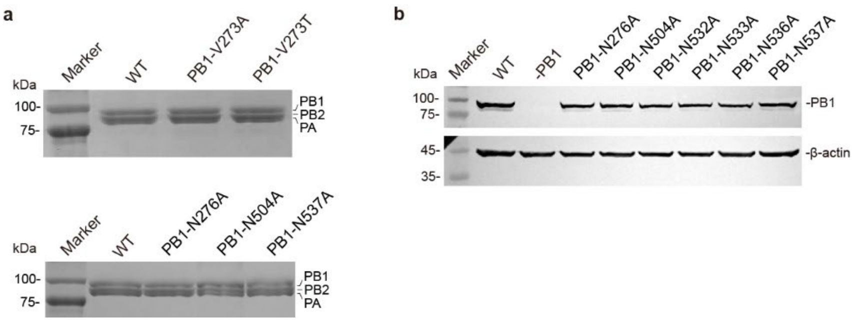


**Fig. S4 The expressions of FluPol$_{H5N1}$. a.** SDS-PAGE analysis was performed on the recombinant proteins of FluPol$_{H5N1}$, including the wild-type, PB1-V273A mutant, PB1-V273T mutant, PB1-N276A mutant, PB1-N504A mutant, and PB1-N537A mutant. The data shown are representative of three independent experiments with similar results. **b.** Western blot analysis was conducted to examine the expression of FluPol$_{H5N1}$ *in vivo*, including the wild-type, PB1-N276A mutant, PB1-N504A mutant, PB1-N532A mutant, PB1-N533A mutant, PB1-N536A mutant and PB1-N537A. The group "-PB1" serves as a negative control. The data shown are representative of three independent experiments with similar results.

**Figure S5**

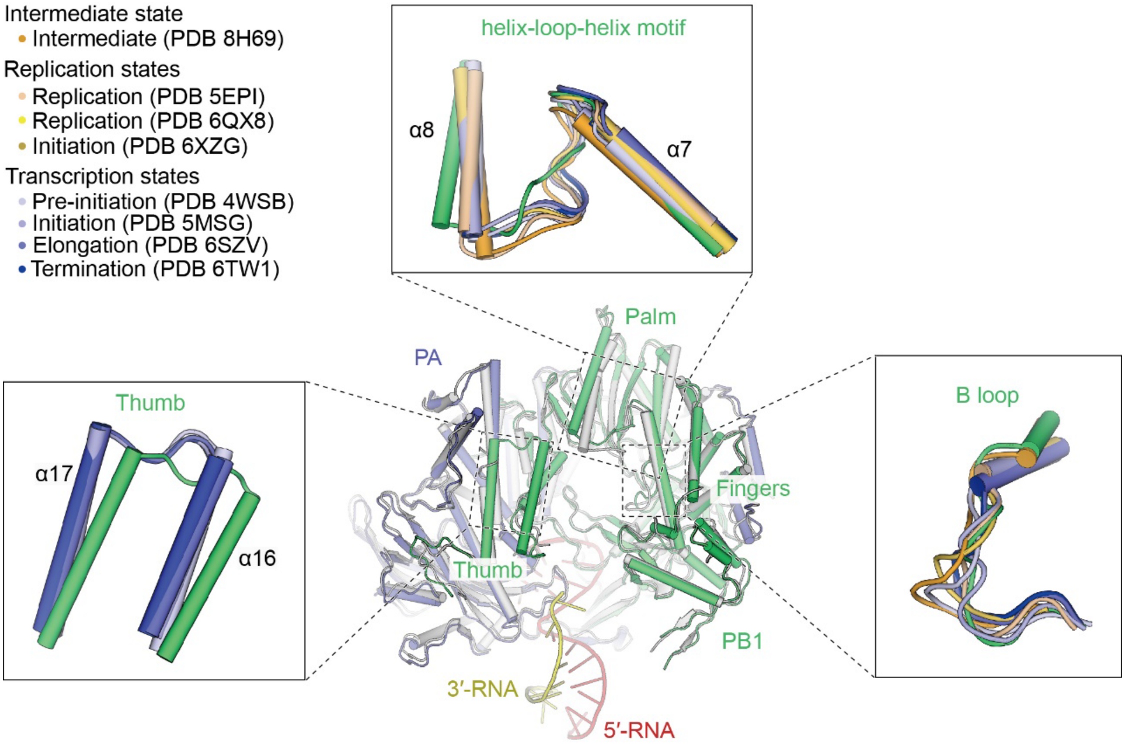


**Fig. S5 The structural features of the FluPol$_{H5N1}$-cRNA structure.** The helix-loop-helix motif, the B loop, the α16 and α17 helices in the FluPol$_{H5N1}$-cRNA structure (PA shown in blue; PB1 shown in green) are compared to the FluPols in intermediate state (orange, PDB 8H69), replication initiation state (yellow orange, PDB 6XZG), replication states (yellow, PDB 6QX8; wheat, PDB 5EPI), transcription pre-initiation state (blue white, PDB 4WSB), transcription initiation state (light blue, PDB 5MSG), transcription elongation state (slate, PDB 6SZV) and transcription termination state (blue, PDB 6TW1).

**Figure S6**

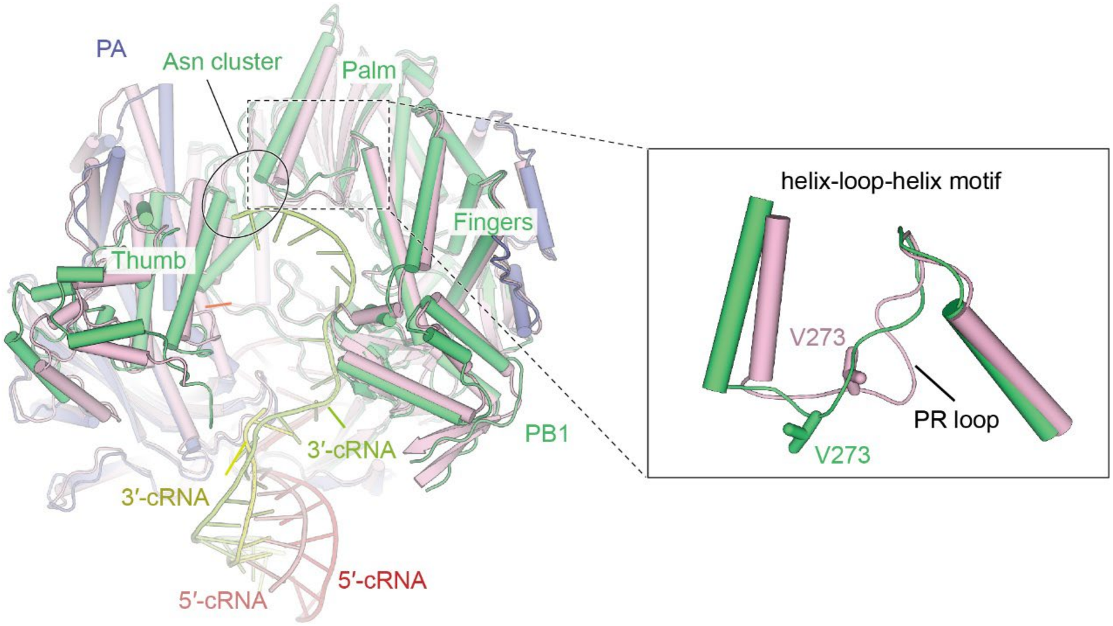


**Fig. S6 Structural comparisons of FluPols bound to the cRNA promoters.** The FluPols binds to the cRNA promoters predicted by the Alphafold 3 server (core region shown in pink, 5′-cRNA shown in salmon, 3′-cRNA shown in limon) is not identical to that determined by the cryo-electron microscopy (PA shown in slate, PB1 shown in green, 5′-cRNA shown in red, 3′-cRNA shown in yellow), particularly the conformation of the residue Val273 in the PR loop, which is essential for priming and realignment process during initiation of vRNA synthesis.

**Figure S7**

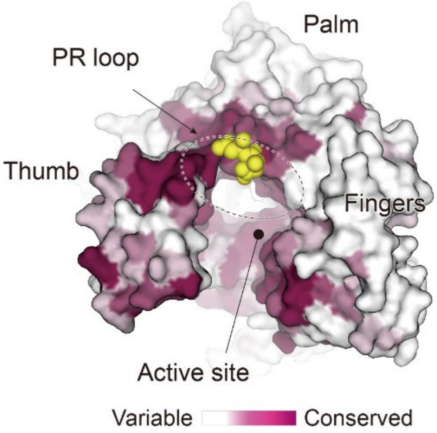


**Fig. S7 The PR loop is highly conserved across the influenza polymerases.** The PB1 subunit of FluPol was binned from white (most variable) to purple (most conserved) using the ConSurf server. The PR loop was shown in the sphere (yellow).

**Table S1 Oligonucleotides**

| Name | Source | Sequence (5′-3′) |
|---|---|---|
| cRNA (45-mer) | Home-made | AGCAAAAGCAGGGUGUUUAAGUU CAAAAACACCCUUGUUUCUACU |
| Capped RNA | Trilink | $m^7$GpppGAAUGCUAUAAUAGC |
| 5′-vRNA (16-mer) | Takara | AGUAGAAACAAGGGUG |
| 3′-vRNA (15-mer) | Takara | CACCCUGCUUUUGCU |
| 5′-cRNA (15-mer) | Takara | AGCAAAAGCAGGGUG |
| 3′-cRNA (16-mer) | Takara | CACCCUUGUUUCUACU |
| 5′-vRNA (18-mer) | BGI | AGUAGUAACAAGGAGUUU |
| 3′-vRNA (17-mer) | BGI | AAACUCCUGCUUUUGCU |
| 5′-cRNA (17-mer) | BGI | AGCAAAAGCAGGGUGUU |
| 3′-cRNA (18-mer) | BGI | AACACCCUUGUUUCUACU |
| mRNA and cRNA primers | BGI | TCCAGTATGGTTTTGATTTCCG |
| vRNA primer | BGI | TGGACTAGTG-GGAGCATCAT |
| 5S rRNA primer | BGI | TCCCAGGCGGTCTCCCATCC |
